# Insights into the microbiota of Asian seabass (*Lates calcarifer*) with tenacibaculosis symptoms and description of *sp. nov. Tenacibaculum singaporense*

**DOI:** 10.1101/472001

**Authors:** Sou Miyake, Melissa Soh, Muhamad Nursyafiq Azman, Si Yan Ngoh, László Orbán, Henning Seedorf

## Abstract

Outbreaks of diseases in farmed fish remain a recurring problem despite the development of vaccines and improved hygiene standards on aquaculture farms. One commonly observed bacterial disease in tropical aquaculture of the South-East Asian region is tenacibaculosis, which is attributed to members of the Bacteroidetes genus *Tenacibaculum*, most notably *T. maritimum*. The impact of tenacibaculosis on fish microbiota remains poorly understood. In this study, we analysed the microbiota of different tissue types of commercially reared Asian seabass (*Lates calcarifer*) that showed symptoms of tenacibaculosis and compared the microbial communities to those of healthy and experimentally infected fish that were exposed to diseased farm fish. The microbiota of diseased farm fish was dominated by Proteobacteria (relative abundance±standard deviation, 74.5%±22.8%) and Bacteroidetes (18.07%±21.7%), the latter mainly comprised by a high abundance of *Tenacibaculum* species (17.6%±20.7%). In healthy seabass Proteobacteria had also highest relative abundance (48.04%±0.02%), but Firmicutes (34.2%±0.02%) and Fusobacteria (12.0%±0.03%) were the next two major constituents. Experimentally infected fish developed lesions characteristic for tenacibaculosis, but the microbiota was primarily dominated by Proteobacteria (90.4%±0.2%) and Firmicutes (6.2%±0.1%). The relative abundance of *Tenacibaculum* species in experimentally infected fish was significantly lower than in the commercially reared diseased fish and revealed a higher prevalence of different *Tenacibaculum* species. One strain was isolated and is described here as *sp. nov. Tenacibaculum singaporense* TLL-A1^T^ (=DSM 106434^T^, KCTC 62393^T^). The genome of *T. singaporense* was sequenced and compared to those of *T. maritimum* DSM 17995^T^ and the newly sequenced *T. mesophilum* DSM 13764^T^.

**Importance:** Fish production from aquaculture facility has become a major source of protein for human consumption and is expected to further grow to meet the growing demands. Devastating fish diseases, such as tenacibaculosis, can eradicate entire stocks of aquaculture fish in a short time and pose a serious threat to individual fish farmers and overall fish production. Understanding the disease processes and the individual microbial players involved has the potential to develop methods to prevent or mitigate infections on aquaculture farms. This study provides important insights into the microbial ecology of tenacibaculosis from an aquaculture facility in Singapore and highlights the complexity of this fish disease at two different disease stages. Furthermore, the isolation of a novel *Tenacibaculum* species and comparative genome analysis of three different *Tenacibaculum* species enhance our view of this economically and environmentally important bacterial genus.

## Introduction

The importance of aquaculture fish as food stock has grown rapidly in recent decades as the amount of wild-caught fish plateaued, while the human world population continues to grow (1, 2). Several technological advances have helped to operate aquaculture facilities in a more sustainable way, both environmentally and economically. For example, different types of water treatments have improved hygiene standards on aquaculture farms and the development of vaccines reduced the number of disease outbreaks in crowded and often genetically uniform farm fish populations (3, 4). Nonetheless, fish disease outbreaks still occur in high frequency. One notable disease example is tenacibaculosis. The disease is characterized by lesions on body surfaces and causes high mortality rates of fish at aquaculture farms (5). The etiological agent of tenacibaculosis was originally isolated from black sea bream (*Acanthopagrus schlegeli*) and was later identified to be the Gram-negative bacterium *Tenacibaculum maritimum* (6).

Since the initial discovery of *T. maritimum*, several other *Tenacibaculum* species have been isolated from different sources around the world, which led to an expansion of the genus to more than 30 different species (5, 7-14). Many of the newly isolated species have been found in association with other marine hosts, such as fish and their eggs (e.g. *T. soleae, T. ovolyticum*) (13, 15), sponges (*T. mesophilum*) (15) and algae *(T. amylolyticum)* (15), but there are also reports that *Tenacibaculum* species may be reaching high relative abundances in water samples (8, 12, 16). However, a comprehensive global survey of the genus *Tenacibaculum* has not been performed thus far. Therefore, certain regions of the world may currently be underrepresented, such as aquaculture facilities in Southeast Asia, despite their significance for the region. It is currently therefore difficult to assess if some of the *Tenacibaculum* species are specific to a host or a geographic region (as some of the species epithets may also suggest (8, 9)).

While diagnostic tools for the detection of *T. maritimum* and other *Tenacibaculum* species have been developed (17), it remains relatively poorly understood how the disease process progresses mechanistically at molecular level. Some indications come from the recently sequenced genome of *T. maritimum*, which revealed that the genome encodes a type IX secretion system (18), that has been discovered only recently and have thus far only been detected in the phylum Bacteroidetes. This translocation machinery has been shown in other Bacteroidetes species, e.g. the human oral pathogen *Porphyromonas gingivalis*, to be involved in virulence factor secretion, but could also be required for gliding motility (19). However, exact functions of the type IX secretion system and prevalence in *Tenacibaculum* species as well of how it contributes to the infection process are currently not fully understood.

Another aspect that has received relatively little attention is the involvement of the commensal fish microbiota in the infection process. It has been shown that animals harbour complex microbial communities on their body surfaces and in the intestinal tract (20-23). These microorganisms not only impact on host physiology, but they may also act as barrier that prevents colonization and infection of pathogenic microorganisms. It is currently not well established how the composition of the fish microbiota (in different tissue types of the fish) changes during tenacibaculosis under non-laboratory conditions, e.g. in fish on aquaculture farms, and how different tissue types are affected.

Here we provide insights into the microbiota of different tissue types of Asian seabass (*Lates calcarifer*) that are affected by tenacibaculosis. Specifically, we looked into the microbiome of four different organ types (gut, skin, kidney and brain) from tenacibaculosis-infected farmed fish and experimentally infected fish and compared them to those of healthy fish. Furthermore, we describe a novel *Tenacibaculum* species, *Tenacibaculum singaporense* that was isolated from experimentally infected fish. *T. singaporense* is characterized, its genome is sequenced and assembled, and the genome annotation compared to *T. maritimum* and the newly sequenced *T. mesophilum* genomes. This study provides novel insights into the microbial ecology of tenacibaculosis and the genomic variability of the genus *Tenacibaculum*.

## Results

### Disease transmission from farm to experimental fish

Tenacibaculosis infected dead fish collected from a local farm were used as inoculum to transmit a tenacibaculosis phenotype to healthy fish in an experimental laboratory tank. Symptoms of the experimentally infected fish included characteristic lesions and necrosis of the various parts of the skin, rotten fins, as well as partially opaque eyes. Additionally, there were clear differences in the weight (two-samples Wilcoxon test, W = 189, p < 0.05) and the total length of the fish (two-samples Wilcoxon test, W = 189, p < 0.05) between the experimentally infected (2.30 ± 0.69g; 56.41 ± 4.56mm) and healthy control tanks (5.83 ± 1.11g; 77.14 ± 4.05mm) at 30 days post-treatment (dpt) when the fishes were sacrificed, indicating abnormal growth is also a trait shared by tenacibaculosis infested Asian seabass. Weight and total length of individual fish are summarized in Table S1.

### Adjustment of sequencing approach and sequencing results

Initially, the primer pair 515FB-806RB, which is widely used as part of the earth microbiome project, was used to amplify and sequence the samples of this study. It was found, however, that 86.2 ± 25.9% of the sequencing reads were assigned to the 18S rRNA gene of the host *Lates calcarifer*. The V1-V2 region of 16S RNA gene (27f-338r) was therefore tested and no detectable amplification of host DNA was found. Internal organs aside from gut (i.e., brain, kidney, head kidney and spleen) could not be sampled from the diseased farmed fish due to severe degradation. Additionally, many sampled tissue types did not yield sufficient bacterial DNA for amplicon sequencing, especially from the healthy fish. Only gut samples could be sequenced from all three treatments (diseased farm, experimentally infected and healthy control fish), while skin samples worked only from the infected fish (i.e. diseased farm and experimentally infected fish). Although some kidney and brain samples produced results from experimentally infected fish, this was not the case for head kidney and spleen which did not yield significant bacterial DNA in neither experimentally infected nor healthy control samples (see Table S1 for a summary of the samples that amplicon sequencing could be conducted and their most abundant phyla). In total, 51 samples were successfully sequenced, yielding 6,360,213 high-quality, non-chimeric sequences across all samples (Table S1). In total, 214,720 bacterial operational taxonomic units (OTUs) clustered at 97% pairwise sequence identity were identified.

### Richness and alpha diversity of bacterial communities

Overall, the mean observed number of OTUs was significantly higher in the control samples (5,754) compared to diseased farm fish (5,490) or experimentally infected fish (3,667), although both diseased farm fish and experimentally infected fish displayed great variation amongst individuals (SD=1,437, 2,166 and 2,390, respectively) (Kruskal-Wallis rank sum test, *χ*^2^ = 12.03, *p* < 0.05) (Table S1 for individual alpha diversity values). In terms of different tissue types, higher number of OTUs were found in the gut (4,991 ± 2,322) than skin (4,787 ± 1,950), kidney (3,508 ± 3,378) and brain (2,568 ± 932), although these differences were not statistically significant (Kruskal-Wallis rank sum test, *χ*^2^ = 7.39, *p* = 0.06). A similar pattern was also detected for the Chao1 index (Kruskal-Wallis rank sum test, *χ*^2^ = 8.50, *p* < 0.05). The clearest difference between the treatments was detected in the Shannon index (Figure 1). The control fish showed a significantly higher diversity (6.05 ± 0.10) than both diseased farm fish (2.60 ± 0.57) and experimentally infected fish (2.76 ± 1.21), although a few samples (~3) from experimentally infected fish also possessed similarly high indices (Kruskal-Wallis rank sum test, *χ*^2^ = 17.44, *p* < 0.05).

**Figure 1.**
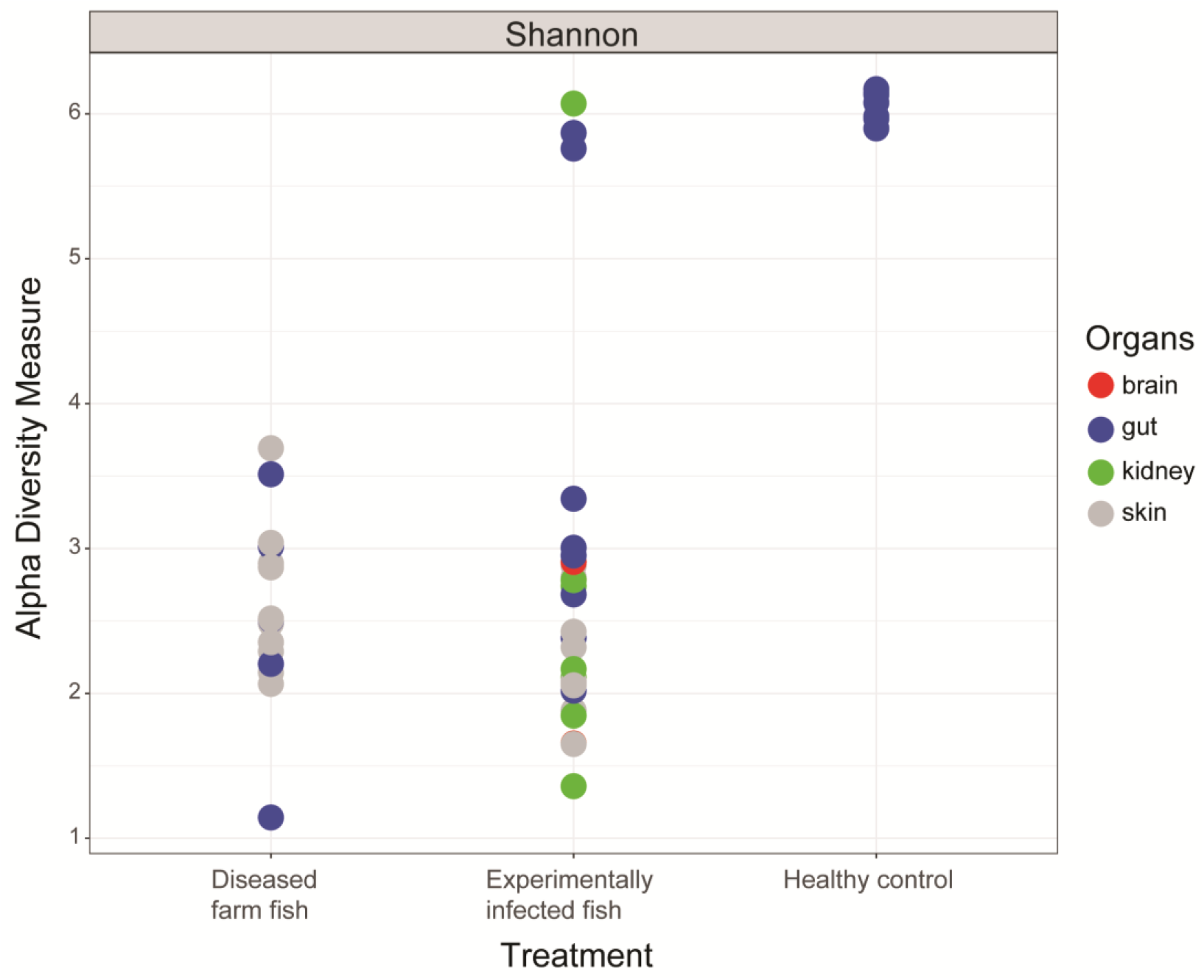
Comparative alpha-diversity analysis of healthy and infected fish. Shannon Index of ‘diseased farm fish’, ‘experimentally infected’ and ‘healthy control’ fish microbiota from various tissue types. Coloration indicates the four different tissue types (brain, gut, kidney and skin) from which microbiota was analysed.

### Taxonomic composition of microbiota in healthy and diseased fish

An in-depth taxonomic analysis indicated that strong differences between the three different groups could be observed at the phylum level. Microbiota samples from the gut – the only organ from which amplification was consistently possible across treatments – of control fish contained higher proportion of Proteobacteria (48.7 ± 2.7 %) and Firmicutes (33.3 ± 0.03%), while diseased farm fish and experimentally infected fish were mostly dominated by Proteobacteria (87.0 ± 15.2% and 83.1 ± 21.0% respectively; see also Figure 2A for details). At the genus level, the highly abundant Proteobacteria in the control fish gut was mostly *Photobacterium* (19.4 ± 3.1%), while the dominant Proteobacteria in diseased farm fish and experimentally infected fish were *Vibrio* (57.3 ± 22.7%) and *Photobacterium* (15.8 ± 11.7%), or *Vibrio* (31.4 ± 28.7%) and *Cohaesibacter* (16.2 ± 24.8%), respectively (Figure 2B; see Table S2A and S2B for full taxonomic abundance at phylum and genus level).

**Figure 2.**
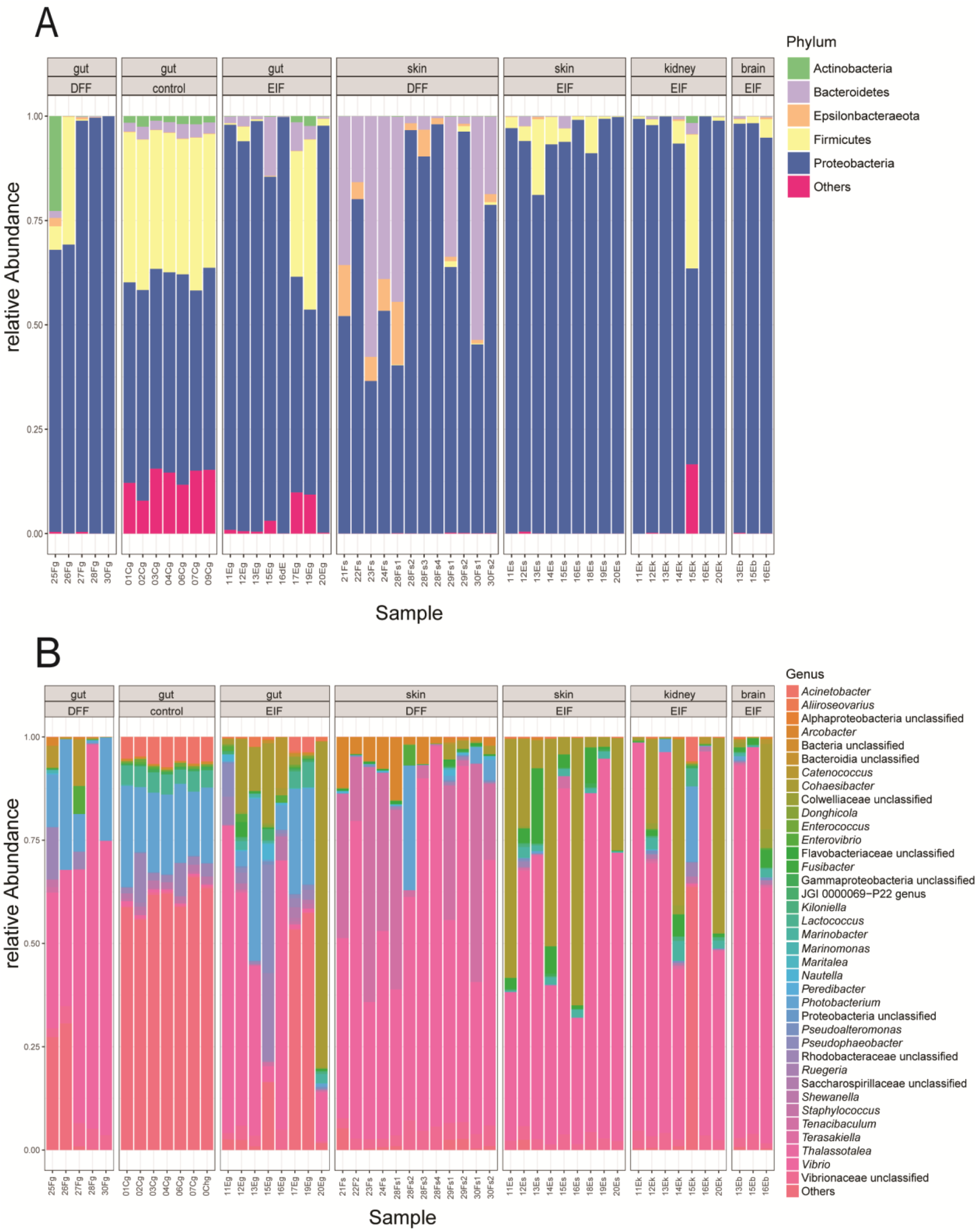
Taxonomic composition of healthy, diseased and experimentally infected fish at different taxonomic levels. A) Phylum-level taxonomy composition of fish microbiota, split by different organ types and condition (‘control’, ‘healthy’ and ‘diseased’). B) Genus-level taxonomy composition of fish microbiota, split by different organ types and condition (‘control’, healthy and diseased). Genera that have a relative abundance of less than 0.01% across all samples are classified as others. Labels: DFF – diseased farm fish; EIF – experimentally infected fish.

The skin microbiome of diseased farm fish and experimentally infected fish microbiome were dominated by Proteobacteria and Bacteroidetes (69.3 ± 22.4%, 25.4 ± 20.3%); or Proteobacteria (94.3 ± 5.5%), respectively. At genus level, the differences became more apparent, with diseased farm fish skin dominated by *Vibrio* (58.0 ± 20.4%) and *Tenacibaculum* (24.8 ± 19.9%), while experimentally infected fish skins were dominated by *Vibrio* (61.7 ± 21.1%) and *Cohaesibacter* (26.3 ± 23.8%). Interestingly, the abundance of *Tenacibaculum* in the experimentally infected fish skin was relatively low (0.4 ± 0.9%). Amplification of microbial 16S rRNA genes was not possible from skin samples of healthy fish.

Amplicons of microbial 16S rRNA genes were also obtained for brain and kidney samples from experimentally infected fish. Both were dominated by *Vibrio* (Proteobacteria; 58.8 ± 31.0% and 61.2 ± 32.1% respectively), although half of brain samples did not contain sufficient bacterial DNA for PCR. The abundance of *Tenacibaculum* was extremely low for both tissue types (0.7 ± 0.8% and 0.1 ± 0.2%, respectively).

### Beta-diversity analysis reveals differences between diseased and healthy fish

The apparent difference in the taxonomic composition between the treatments were further confirmed by beta-diversity analysis. Principal coordinates analysis (PCoA) of Bray-Curtis dissimilarities was performed to determine if differences in microbiota structure between samples from healthy control, experimentally infected and diseased farm fish exist. Although some experimentally infected fish samples clustered together with control samples, most samples clustered according to different treatments, and to less extent, organs (Figure 3). Both axis 1 (representing 33.3% of variation) and 2 (20.7%) separated the treatments. Additionally, an ordination plot of the OTUs at 97% cutoff show distribution of the major OTUs amongst the axis – the three major phyla siding with the three experimental conditions (Figure S1). The difference in the microbiome between the treatments were statistically supported by ANOSIM (*P* < 0.05), as well as the probabilistic modeling to cluster microbial communities into metacommunities by Dirichlet Multinomial Mixtures method (24). The optimum clustering of the samples was three, which can be best explained by the sample treatments.

**Figure 3.**
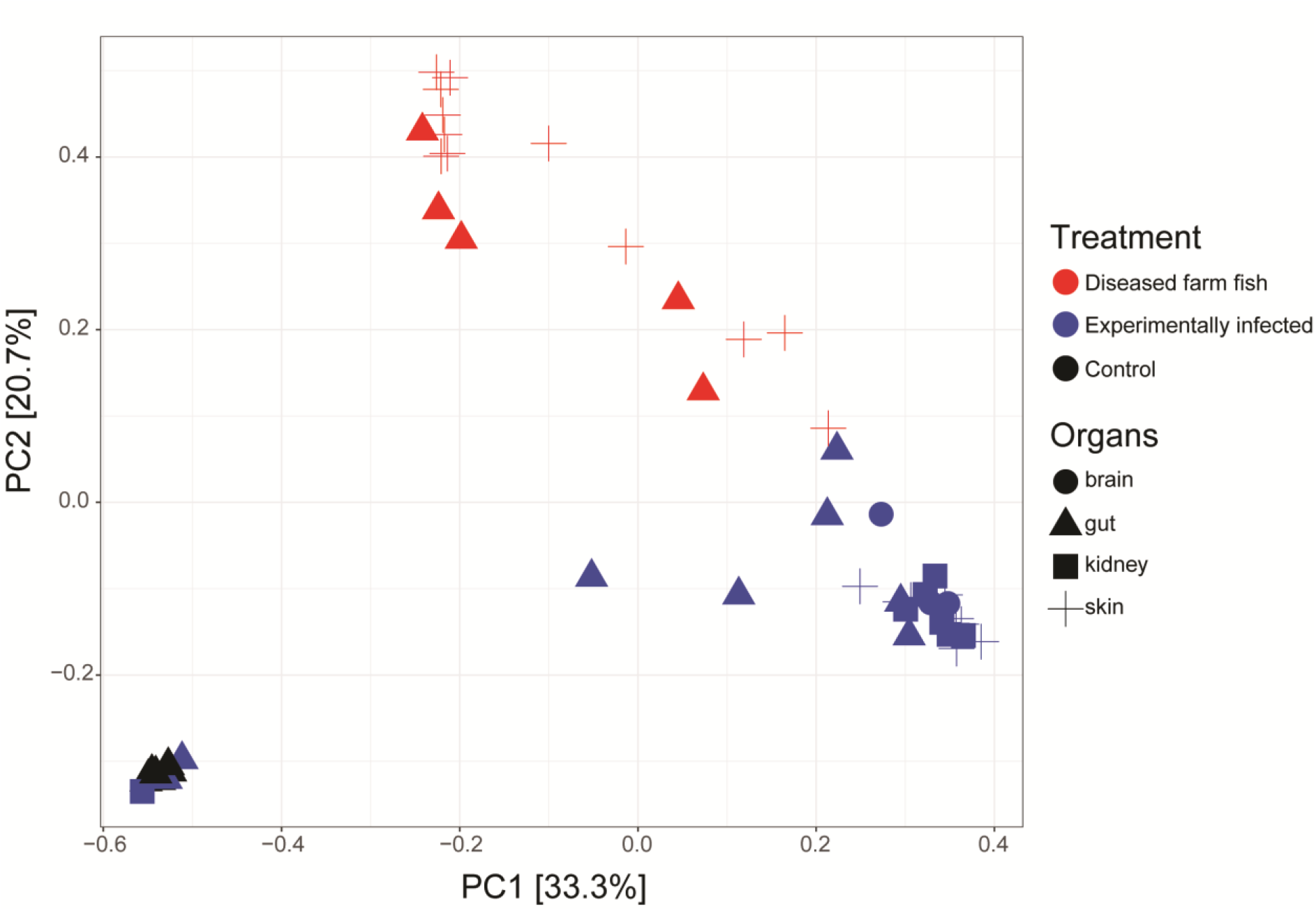
Effects of tenacibaculosis infection on the fish microbiota community structure. Principal coordinate analysis (PCoA) based on Bray-Curtis dissimilarity distances is shown.

Given the strong clustering based on the treatment, we further identified the OTUs responsible for this via LEfSe (Table S3). In total, 365 OTUs were identified to be significantly influencing the clustering, with 43, 32 and 290 OTUs enriched in experimentally infected fish, diseased farm fish and healthy control fish samples, respectively. The experimentally infected fish samples were strongly driven by *Vibrio* OTUs, with five out of the seven most abundant OTUs (OTUs with overall relative abundance > 1%) classified as *Vibrio*, while the other two were *Cohaesibacter*. Although *Vibrio* OTUs were also prominent driver in diseased farm fish samples (four out of seven OTUs over total relative abundance >1% were *Vibrio*), *Tenacibaculum*, *Photobacterium* and *Arcobacter* were also identified as significant discriminants. *Tenacibaculum* OTU in particular was highly abundant (8.9% of total abundance). In contrast, all of OTUs enriched in healthy control samples were below 1% in total relative abundance, perhaps owing to the lower number of samples. Nevertheless, OTUs enriched in them were more diverse, consisting of unclassified Rhodobacteraceae, *Photobacterium, Paracoccus, Lactococcus*, unclassified Fusobacteriales, *Peptostreptococcus, Anthococcus* and *Leuconostoc*.

### Prevalence of *Tenacibaculum* OTUs in different treatments

The five most prevalent *Tenacibaculum* OTUs were picked out from each treatment (diseased farm, experimentally infected and healthy control fish) and placed into the 16S rRNA phylogenetic tree (Figure 4). This revealed that abundant OTUs between the treatments differed. Diseased farm fish possessed diverse *Tenacibaculum* OTUs, with those closest to *T. maritimum* (16.99% of total abundance), *T. lutimaris* (0.61%), *T. skagerrakense* (0.18%), *T. litopenaei* (0.09%) and *T. litoreum* (0.04%). For experimentally infected fish, the three most abundant OTUs where phylogenetically related to *T. singaporense* DSM 106434 (0.191, 0.004, 0.002%), while the other two were closest to *T. mesophilum* (0.124%) and *T. maritimum* (0.003%). The healthy control fish had the lowest relative abundance of *Tenacibaculum* OTUs, with those related to *T. singaporense* (0.01%), *T. dicentrarchi* (0.006%), *T. maritimum* (0.003%) and *T. finnmarkense* (0.001%) being the most abundant.

**Figure 4.**
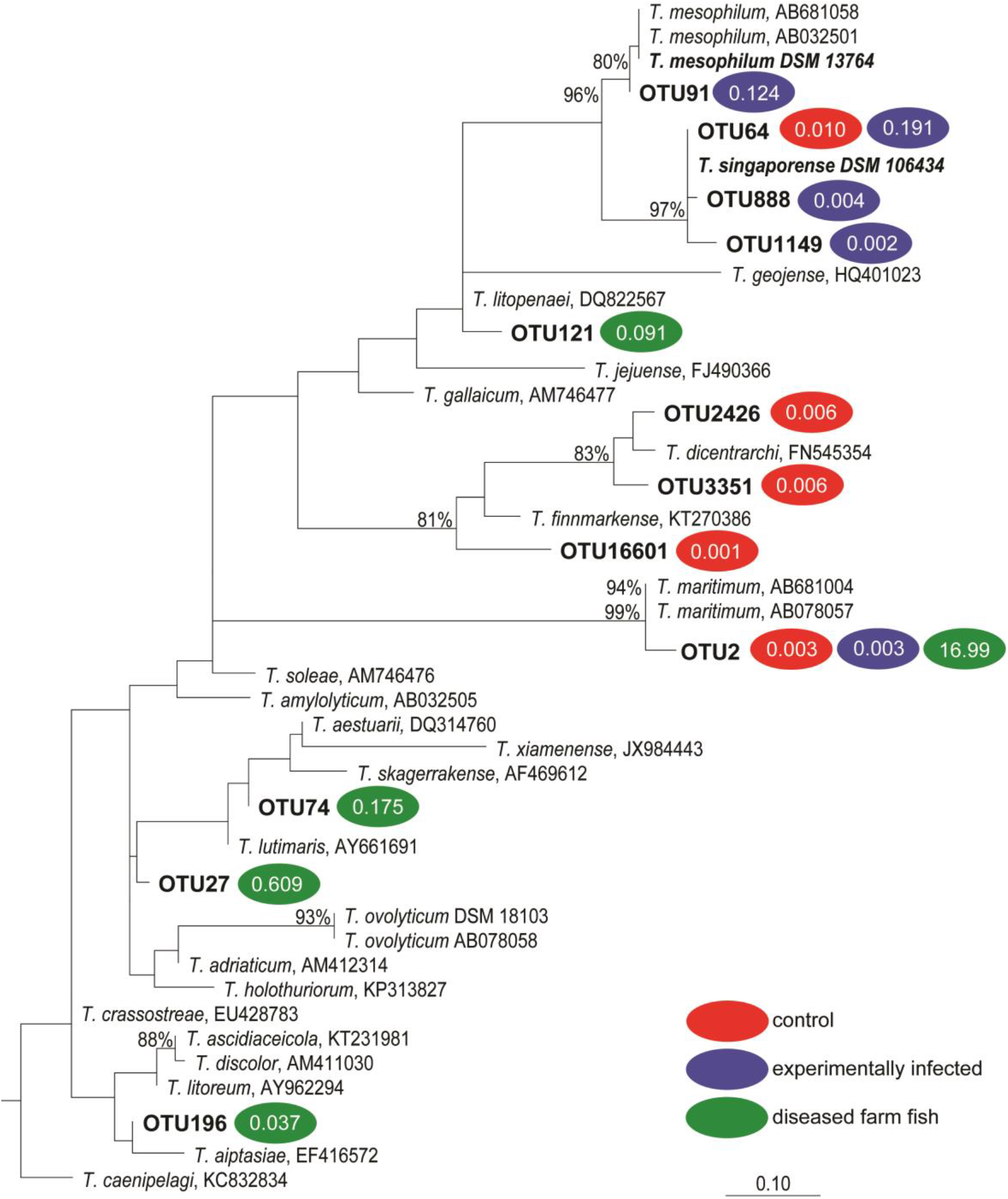
RaxML 16S rRNA gene phylogeny of major *Tenacibaculum* OTUs. The sequences were SINA-aligned (Pruesse et al., 2012) and the tree was constructed in ARB. *T. singaporense* in bold was isolated in this study. The coloured semi-circle denotes the top five most abundant *Tenacibaculum* OTUs from negative control (red), experimentally infected fish (blue) and diseased farm fish (green) with relative abundance noted inside. Bootstrap values >70% are indicated.

**Figure 5.**
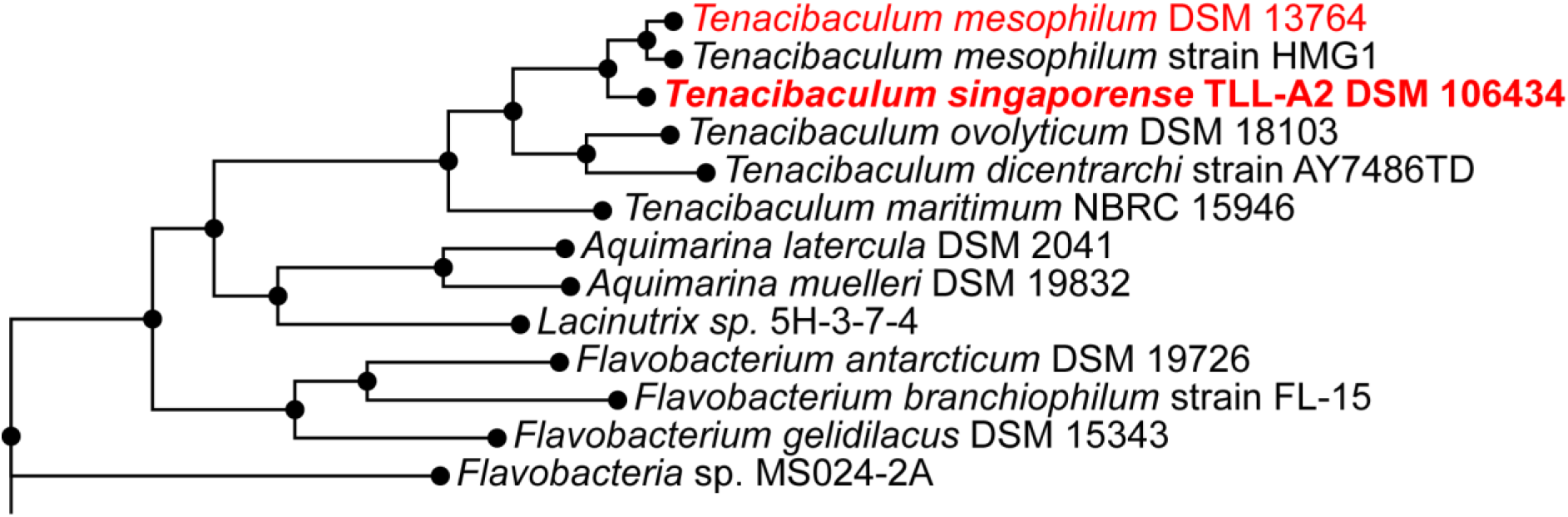
Whole-genome phylogeny of major *Tenacibaculum* genomes. The tree was constructed in PATRIC with RaxML algorithm. Those in red are isolates collected for this study.

### Isolation and characterization of *Tenacibaculum singaporense* DSM 106434

Experimentally infected fish skin displayed lesions characteristic for tenacibaculosis, but the relative abundance of *T. maritimum* in the analyzed samples was very low (0.4 ± 0.9%, compared to 24.8 ± 19.9% in diseased farmed fish skin). However, the analysis revealed the presence of other *Tenacibaculum* OTUs in the experimentally infected fish that could contribute to the disease phenotype. We therefore aimed to isolate the bacteria present in the skin lesion of experimentally infected fish, and one of the obtained isolates showed high sequence identity to the *Tenacibaculum* OTUs with the higher relative abundance. The strain was named *Tenacibaculum singaporense* due to the geographical origin of the isolate (see below for more details on species description). Analysis of the 16S rRNA gene and whole genome phylogeny both showed that *T. singaporense* DSM 106434 is most closely related to *T. mesophilum*, and is phylogenetically distinct from *T. maritimum* that is associated with tenacibaculosis (Figures 4 and 5). The 16S rRNA gene of *T. singaporense* shares 94.5% sequence identity with *T. maritimum* and 98.7% with *T. mesophilum*. Given the similarity of *T. singaporense* and *T. mesophilum* at 16S rRNA gene level, genome-wide average nucleotide identity (ANI) was used to determine the species delineation (**25**). Two-way ANI from 9,697 fragments were 92.07%, well below 95-96% threshold used for species delineation. This was supported by genome sequence-based delineation (GGDC) (**26**), that showed that the probability that DNA-DNA hybridization would be over 70% (i.e., same species) is a mere 9.5% (via logistic regression).

### Phenotypic characterisation of *T. singaporense* DSM 106434

Analyses of general phenotypic features and growth characteristics, including temperature range and colony morphology, of *T. singaporense* DSM 106434 were performed in comparison with *T. adriaticum* DSM 18961, *T. discolour* DSM 18842, *T. maritimum* DSM 17995 and *T. mesophilum* DSM 13764 (for detailed results see Tables S4). The following test were performed by DSMZ: analysis of presence of polar lipids, respiratory quinones, cellular fatty acids, flexirubin. The following lipid groups were identified in the strain: lipid (L), glycolipid (GL), aminolipid (AL), phosphatidylethanolamine (PE) (Figure S2). Analysis of respiratory quinones revealed the presence of menaquinone MK6. Results of cellular fatty acid analysis showed significant proportions of branched-chain and hydroxylated fatty acids (Table S5). Negative test result for the presence of flexirubin indicated that *T. singaporense* does not produce flexirubin.

The ability of *T. singaporense* to utilise casamino acids, N-acetylglucosamine, sucrose, D-ribose, DL-aspartate, L-proline, L-glutamate, hydrolysis of starch, hydrolysis of gelatine, and hydrolysis of chitin was also tested. Casamino acids and hydrolysis of gelatine are the only two of the ten carbon sources found to be used by *T. singaporense*.

Substrate utilisation and substrate derived acid production were tested using API CHE and API 50CH kits (bioMérieux, Craponne France). *T. singaporense* DSM 106434^T^ did not show any reaction with either of the test strips. Results of the API ZYM test kit and the API 20NE test kit are shown in Tables S6 and S7, respectively. Analysis of antibiotic susceptibility indicated that_growth of *T. singaporense* DSM 106434^T^ was not inhibited by oxacillin, gentamycin, amikacin, colistin, pipemidic acid, bacitracin, polymyxin B, kanamycin, neomycin, fosfomycin, and nystatin (see Table S8 for the antibiotic test results).

### Assembly and analysis of the *T. singaporense* and *T. mesophilum* genomes

The genome of the isolated *T. singaporense* DSM 106434^T^ as well as that of the *T. mesophilum* type strain DSM 13764^T^ was sequenced for further analysis. HGAP 4 assembled reads from PacBio RSII sequencing were further error corrected with MiSeq reads using Pilon software, which only corrected for 9 locations totalling 11 bases. Pilon assembly of *T. singaporense* DSM 106434 yielded a single contig for a total assembly size of 3,511,704 base pairs, with G+C content of 32.0%, N50 of 3,511,704 and average coverage of 115x across the genome. No plasmid was identified. The genome has an estimated completeness of 99.66% and contamination of 0.67% based on 548 marker genes conserved in Flavobacteriaceae as identified by checkM. The assembly contained 3,204 coding sequences (CDS); of which 61 were RNAs (8 rRNA and 53 tRNA genes) and 40 repeat regions. This amounted to 1,840 (57.4%) proteins with assigned putative function and 1,364 hypothetical proteins. 2,149 proteins were assigned as FIGfam (see Table S9 for the summary of genome statistics). All annotations are publicly available online under SUB4555753 or in PATRIC ID 104267.16.

Similarly, the Pilon assembly of *T. mesophilum* DSM13764 also yielded a single contig for a total assembly size of 3,344,078 base pairs, with G+C content of 31.8%, N50 of 3,344,078 and an average coverage of 208x across the genome. No plasmid was identified. As was the case for the *T. singaporense* DSM 106434 genome, Pilon only corrected for 8 locations totalling 8 bases. The genome has an estimated completeness of 100% and contamination of 0.61% based on 457 marker genes conserved in Flavobacteriaceae as identified by checkM. The assembly contained 3,044 CDS; of which 66 were RNAs (10 rRNA and 56 tRNA genes) and 52 repeat regions. This amounted to 1,821 (59.8%) proteins with assigned putative function and 1,223 hypothetical proteins. 2,123 proteins were assigned as FIGfam (Table S9). All annotations are publicly available online under SUB4565149 or in PATRIC ID 104268.12.

### Metabolic potential of *T. singaporense* DSM 106434 and *T. mesophilum* DSM13764

The central metabolism of *T. singaporense* and *T. mesophilum* DSM13764 is similar to *T. maritimum* whose metabolic potential has been previously reported (27). Briefly, the genomes possess complete set of genes for glycolytic (Embden-Meyerhof-Parnas) and pentose phosphate pathways, TCA cycle, as well as NADH-dehydrogenase, cytochrome C oxidase and ATP synthase (for the list of enzymes, see Table S10). Additionally, genes for copper-containing nitrogen reductase are present, which share 99% sequence identity at amino acid level between the two genomes.

Carbohydrate Active EnZymes (CAZymes) are involved in the synthesis, breakdown and transport of the carbohydrates. They are classified into glycoside hydrolases (GHs), glycosyl transferases (GTs), polysaccharide lyases (PLs), carbohydrate esterases (CEs), auxiliary activities (AAs) as well as carbohydrate-binding modules (CBMs). Based on predictions, the *T. singaporense* DSM 106434 genome harbors genes for 41 CAZymes, which were identified as 24 GTs, ten GHs, four CEs, one CBM, one PL and one AA (Table S11A). Of 24 GTs found, GT2 was the most abundant (nine), followed by GT4 (five) and GT51 (three). GT2 includes enzymes with chitin synthase, mannosyl and glucosyl transferase activity, which may be involved in the synthesis of chitin and glycosylation of proteins. GT4 are enzymes involved in sucrose and mannose synthesis. GT51 are murein polymerases. In terms of GHs, aside from unclassified GH0 (three), most were lysozyme or chitinase related enzymes (GH23, three; GH18, one; GH73, one).

Similarly, the *T. mesophilum* DSM13764 genome encodes for 36 CAZymes, which can be further broken down into 20 GTs, nine GHs, six CE and one AA (Table S11B). Further GTs breakdown into family followed similar pattern to that of *T. singaporense* DSM 106434, where GT2 was the most common (seven), followed by GH4 (five) and G51 (three). However, unlike in case of *T. singaporense* DSM 106434, there was no clear pattern in GH family classification, with two GH23 and 1 GH113 (both lysozymes), and one each of GH3 (β-glucosidase), GH5 (endo-β-1,4-glucanase/cellulase), GH20 (β-hexosaminidase), GH73 (β-mannanase) represented.

### Predicted virulence potential of *T. singaporense* DSM 106434

Initial RASTtk annotation within PATRIC did not identify any potential pathogenic feature within the *T. singaporense* DSM 106434 genome, and so we conducted BLASTp searches of the *T. singaporense* DSM 106434 genome against Virulence Factor Database (VFDB) core dataset identified 109 significant alignments, which were classified into major virulence factors (Table S12). The most common were toxin formation and iron uptake (both 22.9%), followed by defense (including anti-phagocytosis, immune evasion and antimicrobial activity; 18.3%). Of the toxin formation genes, many were encoding potential hemolysins, which destroy the cell membrane of the host red blood cells. For iron uptake, many were iron transporters and peptide synthase, along with some acinetobactin biosynthesis genes were identified. Adherence (12.8%, mostly in the form of minor curlin and internalin, but also including biofilm formation), stress protein (9.2%), regulation (3.7%) and motility (2.8%) were also found. Less common (and thus classified as others) included secretion system, intracellular survival mechanism and enzymes.

In terms of the organism these factors were found in, many of the hits were from *Pseudomonas aeruginosa* PAO1 (17.4%) and *Haemophilus influenza* Rd KW20 (14.7%). Those originating from the former included iron uptake, adherence and anti-phagocytosis genes, while the latter were toxin, immune evasion and iron uptake genes. We could not determine whether the apparent bias in the origin of virulence factor was from horizontal gene transfer (HGT) as opposed to simply a bias in the database or the genomes of those in the database. Given the possibility of HGTs in acquiring the virulence factors, we further investigated the genome island present in the *T. singaporense* DSM 106434^T^ genome (Figure S3). There were nine major genomic islands identified by at least two methods (Integrated, Island-Path-DIMOB and SIGI-HMM), with the largest one spanning 85,087bp.

### Type IX Secretion System

Secretion systems (SS) are important for pathogens as means of delivering virulence factors. One type 9 secretion system (T9SS) has recently been discovered in the genome of *T. maritimum* (see (19) for details). Homologues of all major components *(porP-porK-porL-porM-porN*) have also been found in *T. singaporense* DSM 106434 and *T. mesophilum* DSM 13764 though not in operon structure as described for some *Bacteroidetes*. The *porP* gene was used as an example to highlight the differences in genomic context between different *Tenacibaculum* species (Figure 6). In *T. maritimum*, four homologues of *porP* genes have been found, with *ompA* and putative adhesion genes directly flanking the gene in all four instances. OmpA is a conserved protein domain common in pathogenic bacteria that may act as porin. In comparison, *T. singaporense* DSM 106434 and *T. mesophilum* DSM13764 have omp16 persecutor and internalin / T1SS secreted agglutinin RTX upstream or downstream of the gene, with *motB* and *tonB* in close proximity as well. MotB is a protein associated with motility, while TonB plays a role in heme utilization, both are often associated with virulence factors.

**Figure 6.**
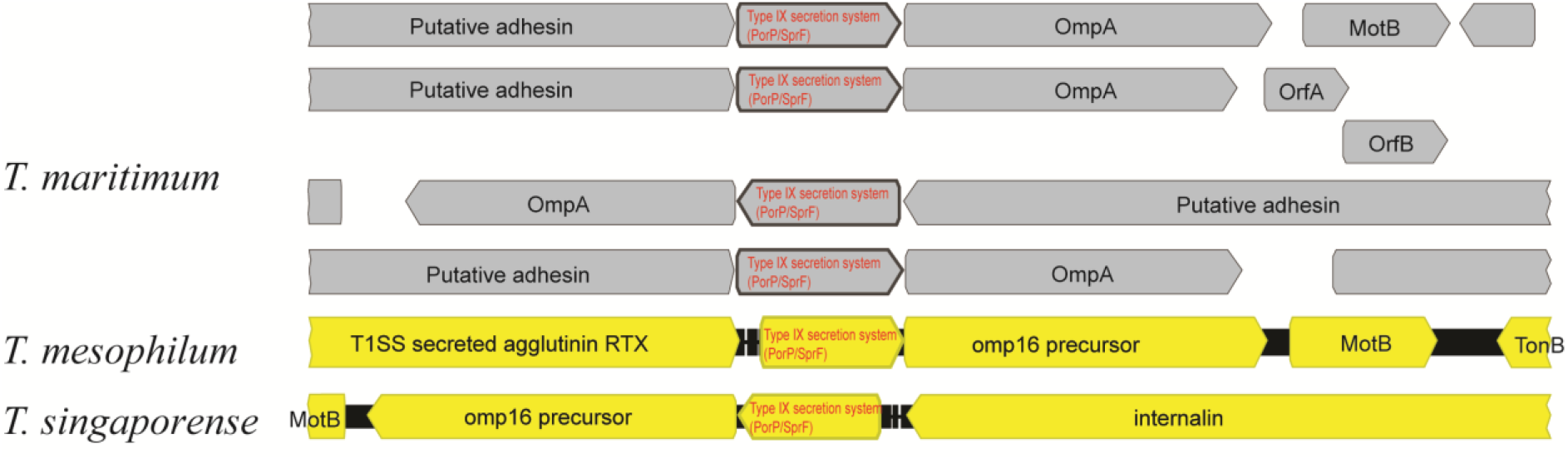
PorP gene (type IX secretion system) and its genomic region annotated in *T. maritimum, T. mesophilum, and T. singaporense* genomes. Four PorP genes were described earlier from the *T. maritimum* genome, while only one was found in *T. mesophilum* and *T. singaporense* genomes each.

### *Comparative analysis of T. singaporense* DSM 106434 *with T. mesophilum* DSM13764 and *T. maritimum genomes*

Circular display representation of the *T. singaporense* DSM 106434 and *T. mesophilum* DSM 13764 genomes against the *T. maritimum* DSM 17995 genome was used to illustrates the similarity between the three different genomes (Figure 7A). An in-depth analysis of the genome content revealed that *T. singaporense* DSM 106434 contained 3,160 proteins, 2,511 COGs and 560 singletons, of which a core of 1,776 COGs were conserved in *T. mesophilum* DSM13764 and *T. maritimum* (Figure 7B). BLASTp analysis of species-specific COGs indicated that most are representing hypothetical proteins with no known function in all three. Interestingly, both *T. singaporense* DSM 106434 and *T. mesophilum* DSM13764 possessed unique TonB-dependent receptor SusC, which is a transport mechanism between outer membrane and periplasm, often found in *Bacteroides*. TonB homologues have been found in vicinity of the *porP* gene, an important component of T9SS, as described earlier. Additionally, *T. singaporense* DSM 106434 possessed homologue of T9SS C-terminal target domain-containing protein from *Tenacibaculum sp*. 4G03.

**Figure 7.**
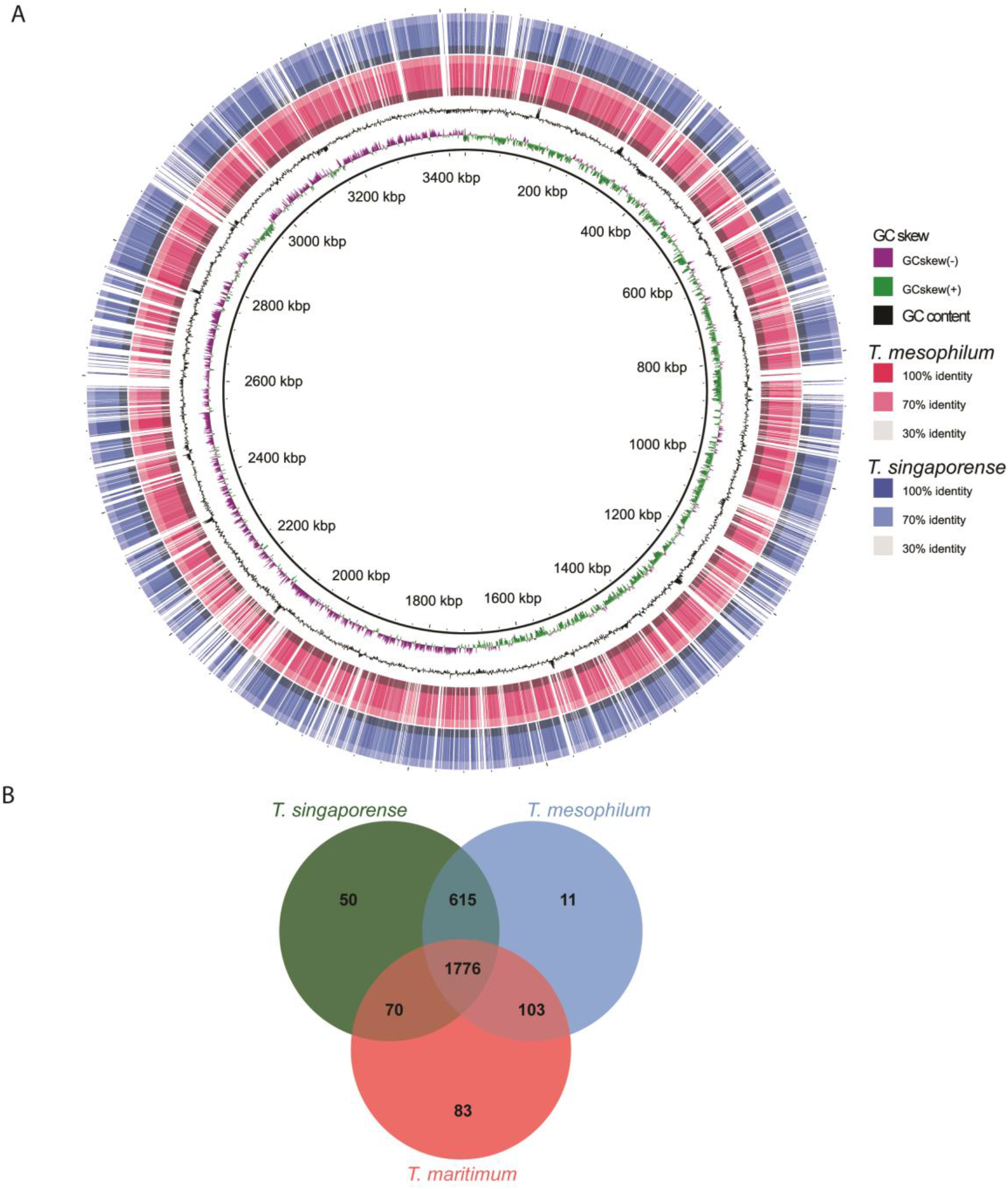
Comparative genome analyses of *Tenacibaculum* species. A) Comparison of *T. singaporense* DSM 106434 and *T. mesophilum* DSM 13764 with *T. maritimum* DSM 17995. BRIG Circular display diagram depicting GC skew, GC content, similarity of *T. mesophilum* (red) and *T. singaporense* (blue) relative to *T. maritimum*. B) Venn diagram of orthologous clusters between *T. singaporense* DSM 106434 isolated in this study, *T. mesophilum* DSM 13764, closest known relative, and *T. maritimum* NCIMB 2154T, prominent tenacibaculosis disease agent.

## Discussion

This study had two main aims. First, the microbiota composition of diseased farm fish with tenacibaculosis symptoms and of healthy Asian seabass was analysed to characterize the disease-specific configuration of the microbiota; Second, a lab-based pathogen challenge was performed by exposing healthy fish to diseased farm fish in order obtain insights into the horizontal transmission process of tenacibaculosis-causing microorganisms. For the generation of data and interpretation of the results it was also important to take the geographic location and the host species for the experiments into account as there is only limited information on the microbial ecology of Asian seabass grown in aquaculture facilities of Singapore and its whole geographical region.

### Technical considerations for the analysis of Asian seabass microbiota

The initial approach of this study to analyse the Asian seabass microbiota relied on the use of primers 515FB and 806RB recommended by the earth microbiome project. These primers have been widely used in many different habitats and are known to have relatively little bias against specific taxa (28). However, it was found in this study that this primer pair mainly amplified a region of the fish ITS1 gene and that only low numbers of amplicon sequencing reads could be assigned to microbial taxa, making it necessary to use a different primer pair. The 27f/338r primer pair used in this study is known to be biased against some taxa, e.g. *Bifidobacteria* (29), but has also been used widely in microbiome studies, it was sufficient for this study to detect the suspected pathogens and to provide insights into the diversity of the Asian seabass microbiome. The increasing interest to analyse fish microbiota may make it necessary to further optimize the required technical approaches. This could include approaches to reduce the concentration and/or amplification of host DNA in the samples or additional modifications of primers, such as the use of blocking primers (30). Considering that fish are the most abundant group of vertebrates with an estimate of more than 34,000 different species (31), it may currently be a difficult task to develop a universal approach that works for many or all species of fish.

### Differences in microbiota composition between healthy fish, diseased farm fish and experimentally infected fish

The alpha- and beta-diversity analysis did reveal strong differences in microbiota composition between healthy and diseased fish, but also between the diseased farm fish and the experimentally infected fish. Although skin is the most obvious site of infection, the difference in the microbiome was pronounced in the internal organs as well. In general, healthy fishes harboured more diverse gut microbial communities in terms of richness and evenness. Some of the detected phyla, e.g. Fusobacteria and Firmicutes, appeared to have only low relative abundances in both groups of the diseased fish, indicating that these phyla might be part of an eubiotic configuration of the healthy Asian seabass microbiota. Fusobacteria and Firmicutes have also been detected in significant numbers in the microbiota of other healthy fishes, including zebrafish (32-34) Atlantic salmon (35) and surgeonfishes (36). It may therefore be worthwhile exploring the presence and abundance of these microorganisms as biomarkers for the fish health status in other species as well. This is corroborated by LEfSe analysis where certain OTUs (that belong to Fusobacteria and Firmicutes) are enriched exclusively in the healthy control samples, as well as correlation analysis which showed that these OTUs are negatively correlated with those that are associated with the disease (i.e. *Tenacibaculum* and *Vibrio*). It remains to be seen if the difference observed here in the gut microbiota between the healthy and diseased fish is a direct effect of pathogenesis, or a secondary effect due to the poor health state of the fish. As Fusobacteria are considered obligate anaerobes (37), it may also be tempting to speculate, whether their absence could be due to disease-induced changes in gut physiology that favour facultative anaerobes or that inhibit strict anaerobes.

In contrast to the gut – where the microbiota was richer in the healthy compared to the diseased – no bacterial marker genes could be amplified from the healthy skin, but from skin of diseased fish, as expected. The lesions of the diseased skin harbored a wide range of bacterial taxa that are potentially pathogenic in nature. Diseased farm fish microbiota revealed a high relative abundance of *Tenacibaculum maritimum* OTUs in samples from fish skin samples. This supports the hypothesis that *T. maritimum* is also the etiological agent of tenacibaculosis in the collected Asian seabass samples as it has also been shown for other fish species in different locations around the world (5). *Photobacterium* and *Vibrio* species also reached high relative abundances (or even higher than *T. maritimum*) in these samples. Both genera are known to harbor also some species that are known fish pathogens (38-40) and may have contributed to the disease progression. However, in the case of *Photobacterium* it should be noted that this genus is also highly abundant in the healthy control fish that do not reveal any obvious signs of disease.

Diseased farm fish and the experimentally infected fish showed similar disease symptoms and a transmission of the potential pathogens, in particular *Tenacibaculum* species, from diseased farm fish to experimentally infected fish appeared to be the likely cause. However, the microbiome composition of experimentally infected fish differed (for most samples) strongly from the diseased farm fish. Experimentally infected fish harboured only a low abundance of *Tenacibaculum* species and were instead mostly dominated by species of the phylum Proteobacteria, most notably *Cohaesibacter* and *Vibrio* species. These observed microbiome differences between the diseased farm fish and the experimentally infected fish raise the question for the underlying cause. Several factors can be envisioned. First, the conditions on the fish farm vary considerably from those in the laboratory. It is known that environmental conditions may affect the composition of the fish microbiota and could therefore also influence transmission of microorganisms from one group to another (41). Second, the microbiota analysis indicates the presence of several other genera with high relative abundance in diseased fish. Therefore, the possibility that some other microorganisms present (i.e. other than *Tenacibaculum* species) may cause a phenotype similar to that of tenacibaculosis (or could have a contributing role) cannot be entirely ruled out. Especially also as tenacibaculosis consist of wide range of symptoms including lesions, frayed fins, tail rot, mouth rot, and can affect multiple fish hosts (42-45). Third, the minimal microbial number of *T. maritimum* cells required to cause tenacibaculosis symptoms in Asian seabass is not well established. The relative abundance of *T. maritimum* in the tissues of experimentally infected fish is nearly 30-fold lower than in diseased farm fish, but depending on threshold (and the absolute numbers) this may still be sufficient to cause the disease. A diagnostic PCR protocol developed specifically for detection of *T. maritimum* (17) did produce positive results for experimentally infected and diseased farm fish skin (data not shown), indicating a substantial colonization level with *T. maritimum*. Fourth, it has to be taken into account that fish were sampled at different stages of the disease and differences in microbiota composition between these two groups may simply mirror different successional stages of the disease. This could also be supported by the finding that some tissue samples, e.g. gut samples of the experimentally infected fish, share higher similarity with the healthy control samples (i.e. not yet affected by the disease due to its early stage), while all diseased farm fish samples clustered with each other, regardless of tissue type, indicating that the infection is in a late systemic stage. Combinations of these four factors or other influences may also have contributed to the microbiome difference, but will require additional investigations.

### *T. singaporense* as a novel representative of the genus *Tenacibaculum* and potential contribution of this species to tenacibaculosis

The analysis of the microbiota of the experimentally infected fish and subsequent cultivation experiments lead to the detection and isolation of a novel *Tenacibaculum* species, *T. singaporense*. This is to the authors’ knowledge the first description of a *Tenacibaculum* species from Singaporean waters. However, there are several other species that have been named after their geographic origin in different regions around the world (8-13, 46). This could indicate that further species from presently not well investigated areas are yet to be discovered. Given the high diversity of *Tenacibaculum* clade, it may also be worth investigating whether certain species or strains are found in higher abundance amongst certain hosts or geographic regions, and if they display any signs of co-diversification, as observed in other host-associated microorganisms (47, 48). Interestingly, this study also reveals the presence of other *Tenacibaculum* OTUs (e.g. sharing high sequence identity with *T. dicentrachi* and *T. skagerrakense*) in the samples of healthy and diseased fish. Finding of *T. singaporense*, together with the finding other *Tenacibaculum* OTUs may indicate that *Tenacibaculum* species could be part of the normal, healthy seabass microbiota. However, it may also indicate that tenacibaculosis could be a multi-factorial disease, with different *Tenacibaculum* species and other bacterial genera potentially influencing the successional pattern of disease progression. In this line it is also interesting to note that it has been shown in mouse experiments that the susceptibility to pathogen invasion could be predicted based on the abundance of closely related species (49), e.g. in the case of seabass it could mean that the presence of *T. singaporense* may increase the susceptibility for a *T. maritimum* invasion, instead of *T. singaporense* being directly pathogenic to fish.

Overall, it may turn out to be highly valuable to obtain better insights into the genome content of *Tenacibaculum* species and the genomic differences between different species. The results of this study indicate that even *Tenacibaculum* species that are only remotely related (based on 16S rRNA sequence identity), such as *T. maritimum* and *T. singaporense*, share a large amount of their genome content, and harbour only relatively few unique COGs, but may also differ in copy numbers of potentially important genes, such as those for type IX secretion system. Undertaking a pan-genome approach for this economically and ecologically important genus and individual species within could help to identify features that allow adaptation to geographic regions and/or to specific hosts.

## Conclusion

In summary, this study provides insights into the microbiota composition of healthy and diseased Asian seabass in a Singaporean aquaculture facility and under laboratory conditions, respectively. The results indicate the presence of a novel *Tenacibaculum* species, *T. singaporense*, which may represent a local relative of the well characterized fish pathogen *T. maritimum* and is closely related to *T. mesophilum*. The ecological significance of *T. singaporense* as well as its contributions to the observed disease phenotype remain currently not well understood and warrant further investigations.

### Description of *sp. nov. Tenacibaculum singaporense*

*Tenacibaculum singaporense*, L. gen. n. *singaporense* from Singapore, in reference to the geographic origin of the isolate.

Gram-negative, oxidase- and catalase positive, strictly aerobe. Grows at temperatures between 20-45°C, in a medium containing 1 to 7% sodium chloride, 30% to 100% Himedia synthetic sea salt medium, and between pH 5 to 9. *T. singaporense* colonies are yellow with an irregular shape and a spreading edge. *T. singaporense* reduces nitrates and utilizes casamino acids and gelatine for growth, but does not produce flexirubin. Its GC-content (inferred from the genome sequence) is 32.01%. *T. singaporense* is TLL-A1 (=DSM 106434^T^, KCTC 62393^T^) was isolated from lesion of diseased Asian seabass *(Lates calcarifer)* in Singapore.

## Material and Methods

### Collection of fish specimens from aquaculture facility

Moribund Asian seabass juveniles (henceforth referred to as ‘fish’ or ‘seabass’) with ca. 5g of body weight with suspected symptoms of tenacibaculosis, having displayed severe skin lesions, and rotten tail and dorsal fins were collected at a local commercial aquaculture facility in Singapore. Images were taken as records. The seabass collected were stored in 80% glycerol at −80°C until usage in the laboratory for microbial isolation and/or the experimental infection experiment.

### Experimental infection experiment

Healthy Asian seabass juveniles of 35 ± 5mm standard length (n=530) were obtained from Marine Aquaculture Centre and quarantined for three days before the start of experiment. Plastic tanks of 200L volume (Toyogo, Japan) were fitted with a 25 watt aquarium heater (Eheim, Germany) to maintain water temperature at 29°C and the fishes were fed to satiation with 80% daily water exchange. Waste water was treated with 10% bleach solution overnight before discharge. Out of the 530 fishes, 470 were subjected to experimental infection and the remaining 60 were kept as untreated control. Moribund seabass collected with suspected tenacibaculosis symptoms (see above) were used as inoculum by adding 15 fishes (either the whole body or body parts with lesions) into the challenge tank. After inoculation, fish were left undisturbed for 10 days until the tenacibaculosis symptoms appeared in the treated tank. Moribund individuals from treated tank were sacrificed humanely by dipping into ice for up to 30 seconds and spiking immediately. All individuals were sacrificed 30 days-post treatment. Gut, infected skin parts (i.e. lesion), brain, kidney, were collected from 10 fish per treatment by sterile scalpels, individually placed in sterile vials and flash frozen by liquid nitrogen. Gut and skin samples were also collected from the diseased farm fish, but other internal organs (brain, kidney, head kidney and spleen) could not be taken as they were partially degraded. The samples were stored at −20°C until processed. Experimental procedures in the laboratory were performed according to the approved IACUC protocol TLL(F)-14–004.

### DNA extraction, library preparation for amplicon sequencing

Samples were transferred to sterile screw-cap tubes containing 0.7g zirconium beads, and 200μl of 20% SDS, 282μl Buffer A, 268μl Buffer PM, 550μl phenol/chloroform/ isoamyl alcohol (25:24:1, pH 8) were added (for details on buffer composition see Rius *et al*. (50). The pelleted cells were subject to phenol-chloroform-based extraction by Rius *et al*. that combined mechanical and chemical lysis of cells. The samples were quantified and diluted to 40 ng/μl per sample and stored in −20 °C for further use.

Extracted DNA was processed according to the Earth Microbiome Project (http://www.earthmicrobiome.org/protocols-and-standards/16s/) as described in Thompson et al (51) except for two points. Firstly, primer pair 27f and 338r spanning hypervariable region V1-3 was used instead of the designed primer pair (515FB and 806RB). Secondly, a dual-indexing strategy was used as described by Fadrosh et al. to reduce the number of barcoded primers required (52). PCR was conducted with QIAGEN Taq MasterMix (CAT NO 201445, QIAGEN, Germany) in triplicates (plus one negative control per sample) under the condition: 94 °C for 3 mins; 35 cycles of 94 °C for 45 secs, 50 °C for 60 °C, 72 °C for 90 secs; and a final elongation at 72 °C for 10 mins. The amplicons were quantified with Quant IT Picogreen (CAT NO P7589, Thermo Fisher Scientific, USA), pooled together in equimolar concentrations and sequenced at Singapore Centre for Environmental Life Sciences Engineering (SCELSE) at Nanyang Technological University using Illumina MiSeq paired-end chemistry (251 × 251bp). The output raw sequences were analysed with MOTHUR v1.39.1 (53) according to the standard MiSeq protocol (54). The sequence pairs were merged, de-multiplexed and quality-filtered accordingly. Visualization of processed data was conducted by R project following tutorials provided in https://joey711.github.io/phyloseq/index.html. Packages used include: ape (55), dplyr (56), ggdendro (57), ggplot2 (58), ggpubr (59), gplots (60), grid (61), gridExtra (62), Heatplus (28), pheatmap (63), phyloseq (64), plyr (64), RColorBrewer (65), reshape2 (66), sfsmisc (67), vegan (68) and viridis (29). Difference in the alpha and beta diversity between treatments / tissue types were tested for significance using Kruskal-Wallis rank sum test (69) and ANOSIM (70), respectively. Dirichlet Multinomial Mixtures method was used to determine the optimal number of clustering in our data (24), and linear discriminant analysis effect size (LEfSe) was used to determine the statistically significantly enriched OTUs from each treatments (71).

### Isolation and cultivation of *Tenacibaculum singaporense*

Sterile cotton buds were used to swab infected skin of the fish showing tenacibaculosis symptoms from the tank experiment and subsequently transferred to marine agar plates prepared in-house (BD Difco^TM^). The agar plates were incubated at 25°C, and the colonies were diluted and streaked out onto them three times to ensure their purity. The identity of colonies was determined by PCR and sequencing of the 16S rRNA gene (27F/1492R) and a single isolate identified as *Tenacibaculum* was kept for further analysis. The isolate was deposited in Deutsche Sammlung von Mikroorganismen und Zellkulturen (DSMZ, Braunschweig, Germany) under strain number DSM 106434^T^ and with the Korean Collection for Type Cultures (KCTC, Jeongeup, Republic of Korea) under strain number KCTC 62393^T^.

### Cultivation of *Tenacibaculum adriaticum*, *T. discolor*, *T. maritimum*, *T. mesophilum and T. singaporense*

*T. adriaticum* DSM 18961^T^, *T. discolor* DSM 18842^T^, *T. mesophilum* DSM 13764^T^ *and T. maritimum* DSM 17995^T^ were purchased from DSMZ as freeze-dried cultures. *T. discolour* was maintained in Marine Broth 2216 (BD Diagnostics, Durham, North Carolina, USA) and incubated in shaking incubator at 25°C, 180rpm. *T. mesophilum, T. maritimum*, and *T. singaporensis* were maintained in marine broth (BD Difco^TM^) and incubated in shaking incubator at 30°C, 200rpm. *T. adriaticum* was maintained in ½ strength marine broth and incubated in shaking incubator at room temperature, 180rpm. Identity of the strain was confirmed via sequencing of the 16S rRNA gene before and after completion of the characterization experiments and extraction of genomic DNA for whole genome sequencing, respectively.

### Phenotypic characterisation of *Tenacibaculum* species

#### Gram staining

Gram reactivity *T. singaporense* DSM 106434 was tested using the Gram stain reagents by crystal violet, Gram’s iodine solution, and Gram’s safranin solution (Sigma-Aldrich, St. Louis, Missouri, USA) and decoloriser solution comprised of 1:1 ethanol (Fisher Scientific U.K. Limited, Leicester, United Kingdom) to acetone (Merck Specialities Private Limited, Mumbai, India). Protocol used was as described by Sigma-Aldrich Gram Staining Kit.

#### Growth response of *T. singaporense* DSM 106434 to different cultivation conditions

Growth and salinity experiments were conducted with a base medium of 1/5 Luria-Bertani medium (LBM), as recommended (15). For every 1L of Himedia artificial sea water salts broth (modified after (72)), 2g tryptone (Oxoid Ltd., Basingstoke, United Kingdom), 1g yeast extract (Oxoid Ltd., Basingstoke, United Kingdom) were added. Fifteen grams of Bacto agar (BD Diagnostics, Durham, NC, USA) was added to five-fold diluted LBM agar, and pH was adjusted to 7.5 with 5M NaOH (Schedelco, Singapore). The medium was sterilised by autoclaving for 20 mins at 121°C. To obtain above mentioned Himedia artificial sea salts, the following were added per every litre of water: 24.6g of sodium chloride (Merck, Hellerup, Denmark), 0.67g of potassium chloride (Sigma, St. Louis, MO, USA), 1.36g of calcium chloride dihydrate (Merck, Darmstadt, Germany), 3.07g of anhydrous magnesium sulphate (Sigma-Aldrich, Tokyo, Japan), 4.66g of magnesium chloride hexahydrate (Sigma-Aldrich, Munich, Germany), and 0.18g of sodium bicarbonate (Sigma-Aldrich, St. Louis, Missouri, USA). Final pH at 25°C was adjusted to pH7.5 ± 0.05.

Growth response of *T. adriaticum, T. discolor, T. maritimum, T. mesophilum and T. singaporense* DSM 106434 to different salinity levels, synthetic sea water concentrations, and pH values were tested. Growth response to varying salinity was tested in 1/5 LBM broth with varying NaCl concentrations (1, 3, 5, 7 or 10% (w/v) NaCl); to varying synthetic sea water concentrations in 1/5 LBM broth with varying Himedia synthetic sea water concentrations (0, 10, 30, 50, 70, 100% broth); and to different pH levels in 1/5 LBM broth at pH 3, 5, 7, 9, 10. Different pH values were obtained by adding 37% fuming hydrochloric acid (Merck, Darmstadt, Germany) and 4M NaOH (Schedelco, Singapore) as required to 1/5 LBM broth. For all three tests, two serial overnight cultures were grown in 2.5ml medium with respective salinity and pH values. The experiments were conducted in triplicates for each species and each medium composition. Of the second overnight culture, 250μl samples were inoculated into fresh 2.5ml medium (with the respective test condition). On the fifth day from first inoculation, samples were checked for growth. Liquid cultures of *T. mesophilum*, and *T. singaporense* DSM 106434 were incubated at 30°C, shaking at 200 rpm.

The ability of *T. singaporense* DSM 106434^T^ to grow in anaerobic conditions was investigated on solid medium. 1/5 LBM agar plates were placed in an anaerobic chamber for two days to become anaerobic. Five μl live cultures were then inoculated onto centre of each agar and incubated in anaerobic chamber at 24°C.

Growth of *T. singaporense* DSM 106434^T^ at different temperatures was tested by incubating each strain on 1/5 LBM agar plates at 4, 20, 30, 37, 40 and 45°C for up to 6 days. Five μl overnight cultures of each species were used on each agar plate. Three replicates per temperature condition and species were tested.

#### Characterization of enzyme activities, substrate utilization, pigments, respiratory quinones and antibiotic susceptibility of *T. singaporense* DSM 106434^T^

Catalase production was determined using 3% H_2_O_2_ as previously described (73). Oxidase production based on N,N-dimethyl-p-phenylenediamine oxalate and α-naphthol (74) was determined using oxidase test discs from Sigma-Aldrich (Sigma-Aldrich, Bangalore, India). The following tests for enzymatic activity and substrate utilization of *T. singaporense* DSM 106434 were performed by DSMZ: Growth on casaminoacids, N-acetylglucosamine, sucrose, D-ribose, DL-aspartate, L-prolin, L-glutamate, hydrolysis of starch, hydrolysis of gelatine, and hydrolysis of chitin), metabolic traits using API 50CH, API CHE, and API ZYM kits. Analysis of polar lipids, flexirubin test, API 20NE (24 to 48-hour identification of Gram negative non-Enterobacteriaceae), analysis of respiratory quinones, and analysis of cellular fatty acids. Antibiotic susceptibility of *T. singaporense* DSM 106434 to 36 antibiotics was also performed by DSMZ.

#### Extraction of genomic DNA for whole genome sequencing

*T. singaporense* DSM106434 and *T. mesophilum* DSM13764 were grown in liquid medium to an optical density of 1.80 at a wavelength of 600nm. Five-hundred ml culture per isolate were centrifuged (Beckman Coulter rotor JA10, 6000g, 20min, 4°C) to pellet the cells out of the media. Genomic DNA (gDNA) was prepared using two rounds of phenol-chloroform purification (modified after Sambrook et al. (**75**)) and stored in −20°C until further use.

Ten μg of prepared genomic DNA was purified with AMPure XP magnetic beads and quality checked with Nano drop and Qubit fluorometer (Invitrogen, Carlsbad, CA, USA) and subsequently used for PacBio RSII Single Molecule and Illumina MiSeq paired-end (251 × 251bp) sequencing at Singapore Centre for Environmental Life Sciences Engineering (SCELSE) in Nanyang Technological University.

#### Genome assembly

Raw PacBio sequencing reads were *de novo* assembled with SMRT analysis software (via HGAP 4) with standard parameters, except the estimated genome size was set as 3.44Mb according to the *T. maritimum* genome available (27). Reads from MiSeq pair-end sequencing were then used for further error correction using Pilon (76).

As an initial quality check step of the assembly, RNAmmer (77) was used to annotate and verify the RNA genes, of which 16S rRNA gene was further used for phylogenetic tree construction. In addition, CheckM (78) was used to assess genome completeness, percentage contamination, as well as finding out missing single-copy marker genes. Genomes were annotated using RAST online (79) and RASTtk (80) through PATRIC (Snyder et al., 2007). Functional characterization was also performed using PATRIC.

#### Single-gene and whole-genome phylogenetic analyses

16S rRNA gene and whole genome phylogenies were constructed in order to compare the evolutionary placement of *Tenacibaculum* strains with the previously published *Tenacibaculum* sequences. 16S rRNA genes were extracted from the polished genome using RNAmmer as described earlier, aligned by SINA (81) and phylogeny constructed by RAxML (82) within ARB (83). Whole genome phylogeny was constructed in PATRIC using conserved protein sequences via RAxML.

#### Genome-based species delineation

In order to determine if *T. singaporense* DSM 106434 qualifies to be given new species status, two independent methods were used that compared its genome with that of the closest known sequenced type strain, *T. mesophilum* DSM13764, also sequenced in this study. The first approach is based on genome-wide average nucleotide identity (ANI) using reciprocal best hits as described previously (25). Typically, genomes are considered to belong to the same species for ANI values above 95%. The analysis was conducted via online server (84). The second method is genome sequence-based delineation (GGDC) which performs *in silico* DNA-DNA hybridization and determines the probability that two genomes belong to the same species (>70% DNA-DNA hybridization).

#### Genome analysis and metabolic reconstruction

Genome annotation, comparison and metabolic construction, was conducted in PATRIC webserver (85). Presence and absence of major central metabolism genes (i.e. Embden-Meyerhof-Parnas pathway, citric acid cycle, pentose phosphate pathway, NADH-dehydrogenase, cytochrome C, cytochrome C oxidase, ATP synthase genes) were checked in both *T. singaporense* DSM 106434^T^ and *T. mesophilum* genomes. Carbohydrate-active enzymes (CAZymes) were annotated via dbCAN2 meta server (86) and the major class of CAZy checked against the online database (87). Note that only CAZymes identified with >2 methods (out of HMMER (E-Value < 1e-15, coverage > 0.35; http://hmmer.org/) (88), DIAMOND (E-Value < 1e-102) (89) and Hotpep (Frequency > 2.6, Hits > 6) (90) were considered, as recommended by the authors.

In order to determine the potential virulence genes, BLASTp search was conducted on the genome annotation of *T. singaporense* DSM 106434 against Virulence Factors Database (VFDB) core database (A) which consists of experimentally verified virulence factors. Relatively conservative cut-off of E-value <0.01, >70% coverage and >30% identity was used to determine the significant matches. Potential genomic islands were visualized using IslandViewer4 (91). Furthermore, given the significance of type IX secretion system, its major components (porP-porK-porL-porM-porN) were BLAST searched in the genome. Furthermore, since porP was annotated in *T. maritimum* (GenBank ID: LT634361), we used it to align the gene against *T. singaporense* DSM 106434 and *T. mesophilum* genomes in Geneious 8.1 (http://www.geneious.com) (92), and the alignment (including genes up- and downstream) were studied.

#### Comparative genome analyses

In order to visualise the overall differences between *Tenacibaculum* genomes, BLAST Ring Image Generator (BRIG) was used to display the genome contiguity, GC content as well as GC skew and BLAST identity compared to reference *T. maritimum* (strain NCIMB 2154T) (93). Furthermore, OrthoVenn webserver (94) was used for comparisons and annotation of orthologous gene clusters between the three genomes. The unique, shared (between two genomes) and conserved (between all three genomes) Clusters of Orthologous Groups of proteins (COGs) were further analysed using BLAST.

#### Accession numbers

The demultiplexed, pair-matched amplicon sequences are deposited in NCBI SRA under accession number SUB4555782. The assembled *T. singaporense* DSM 106434 and *T. mesophilum* DSM 13764 genomes were deposited in GenBank under accession numbers SUB4555753 and SUB4565149, respectively.

## Acknowledgments

We thank Daniela Moses at SCELSE Singapore for performing the PacBio sequencing, and Amit Anand and Muhammad Khairillah Bin Nanwi at TLL biocomputing for bioinformatics support. This work was funded by Temasek Life Sciences Laboratory.

